# DISGENET: Accelerating Data-Driven Discovery in Disease Genomics and Therapeutic Development

**DOI:** 10.64898/2026.01.05.697749

**Authors:** Janet Piñero, Javier Corvi, Natalia Rykova, Anna Guillem, Amelia Martínez, Jaione Telleria Zufiaur, Ivo Rivetta, Sarah Holmes, Mariandrea Del Boccio, Daniel Shapoval, Macarena De Sarasqueta Szneiderowicz, Felix Slager, Ferran Sanz, Laura I. Furlong

## Abstract

Precision medicine and therapeutic development rely on a comprehensive understanding of genotype–phenotype relationships, yet this information remains fragmented across diverse sources. DISGENET, established over 15 years ago, addresses this challenge by systematically integrating gene–disease, variant–disease, and disease–disease associations from authoritative databases and the literature. This major upgrade expands coverage with chemical and pharmacological annotations and integrates biobank and clinical data. An advanced natural language processing (NLP) pipeline captures emerging evidence with full provenance and contextual details, key to streamlining data-driven insights. DISGENET supports diverse users through multiple tools, including an intuitive web interface, a REST API, an R package, a Cytoscape app, and an AI assistant for natural language queries. Quarterly updates ensure data currency, while a sustainable freemium model provides free academic access and supports ongoing development. DISGENET aims to accelerate data-driven discoveries and advance precision medicine and drug development. The platform is accessible at https://www.disgenet.com.

**Graphical Abstract:** 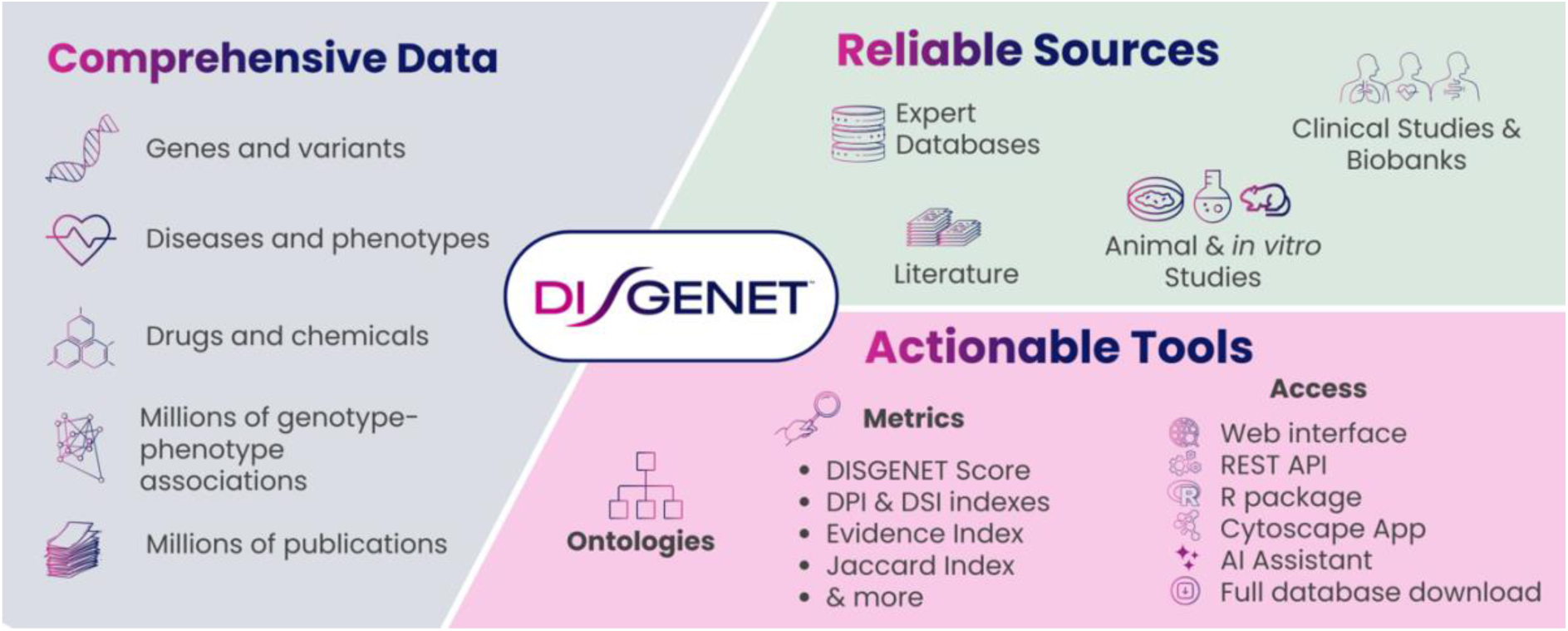

**Highlights:**

- DISGENET accelerates precision medicine via comprehensive genotype–phenotype data
- Access to gene, variant, disease, and drug data in a single platform
- Up-to-date evidence with provenance and full context
- Suite of tools to serve diverse research and clinical communities
- Sustainable freemium model ensures ongoing platform innovation.

## Introduction

Understanding the genetic underpinnings of human disease is a central and enduring challenge in modern biomedicine. The exponential growth and technological diversity of genomic data have paradoxically created an immense yet fragmented landscape of information. This fragmentation, often conceptualized as “data silos,” hinders the efficient translation of raw data into actionable knowledge. Compounding this challenge, a significant volume of new scientific discoveries remains embedded in the text of scientific publications and other textual sources (e.g., clinical trial reports), and these discoveries are incorporated into structured databases at a slow pace.

More than 15 years ago, the original DisGeNET knowledge base was established to directly address these challenges by consolidating information on gene-disease associations (1). This pioneering approach integrated data from specialized, expert-curated databases and automatically mined from the scientific literature, providing a comprehensive view of the genetic etiology of human diseases and phenotypes (2–7). The original platform’s utility and reliability were widely acknowledged by the research community, as evidenced by the continued increase in citations and its recognition as an ELIXIR Recommended Interoperability Resource in 2018^1^. This widespread adoption and long-standing presence in the field underscore DISGENET’s established credibility and the trust it has earned within the scientific community. The resource is not a new tool but a mature, continuously evolving platform that has consistently adapted to community needs and technological advancements. The new DISGENET platform is the latest evolution of this work, building upon its foundational principles while significantly expanding its capabilities to meet modern demands in translational research, especially for precision medicine and therapeutic discovery.

This article presents the new DISGENET platform, which represents a substantial advance over its predecessor in scope, functionality, and sustainability. DISGENET hosts a comprehensive and reliable knowledge base on the molecular mechanisms of human diseases. By combining advanced Natural Language Processing (NLP), semantic technologies, and biomedical data integration, DISGENET streamlines access to high-quality evidence linking genes, proteins, and genomic variants to diseases. This information is extensively annotated with drugs and chemicals, including endogenous and environmental compounds, to provide valuable insights into drug response and environmental effects on disease risk. This upgrade encompasses a massive increase in data content, an expanded suite of access tools designed for diverse user needs, and a novel sustainability model that ensures long-term availability, continuous development, and shorter update cycles. The objective of this publication is to provide a comprehensive description of the complete resource, detailing its scientific and technical improvements, and showcasing its application in translational research. The platform is accessible at https://www.disgenet.com/.

### Data Content and Expansion

DISGENET’s primary mission is to provide the most comprehensive collection of genotype–phenotype associations available. The latest release (v25.3) marks a significant expansion over previous versions, significantly increasing the amount of data and further solidifying the platform’s leading position in integrated genomic knowledge. Specifically, this release features a 1,183%^2^ increase in variant–disease associations (from 369,554 to 4,740,453 VDAs), a 753% increase in variants (from 194,515 to 1,659,055), a 78% increase in gene–disease associations (from 1,134,942 to 2,016,667 GDAs), a 40% increase in genes (from 21,671 to 30,377), and a 27% increase in diseases and phenotypes (from 30,171 to 38,181).

A key enhancement is the integration of information on therapeutic drugs and chemicals, which contextualizes genotype-phenotype associations within a pharmacological and toxicological framework and adds the environmental factor dimension to complex diseases. This new dataset, which, in its current version, covers more than 12,000 drugs and chemicals collated from databases and the scientific literature, was not available in the latest published version of the platform (7).

### The DISGENET data model

A central feature of DISGENET is its ability to aggregate and integrate results from a broad variety of experimental approaches used to uncover genotype-phenotype associations into a single, unified resource. This includes evidence derived from experimental disease models (such as mouse models or *in vitro* cellular systems), biochemical and molecular biology studies, clinical research involving patients (including genome sequencing or GWAS studies on population cohorts), and clinical trials, among other sources.

The gene-disease association (GDA) concept, pioneered by DISGENET, integrates evidence from diverse sources accumulated over years of research into a single association (Figure 1). This design allows recent discoveries about a particular GDA to be contextualized with previous findings, enabling users to assess the association’s novelty or established nature. In addition, chemicals and drugs that influence the GDA are extracted from the textual sources supporting the association. Users can explore the underlying evidence supporting each association, and various annotations and metrics are provided to streamline data prioritization and interpretation.

**Figure 1:**
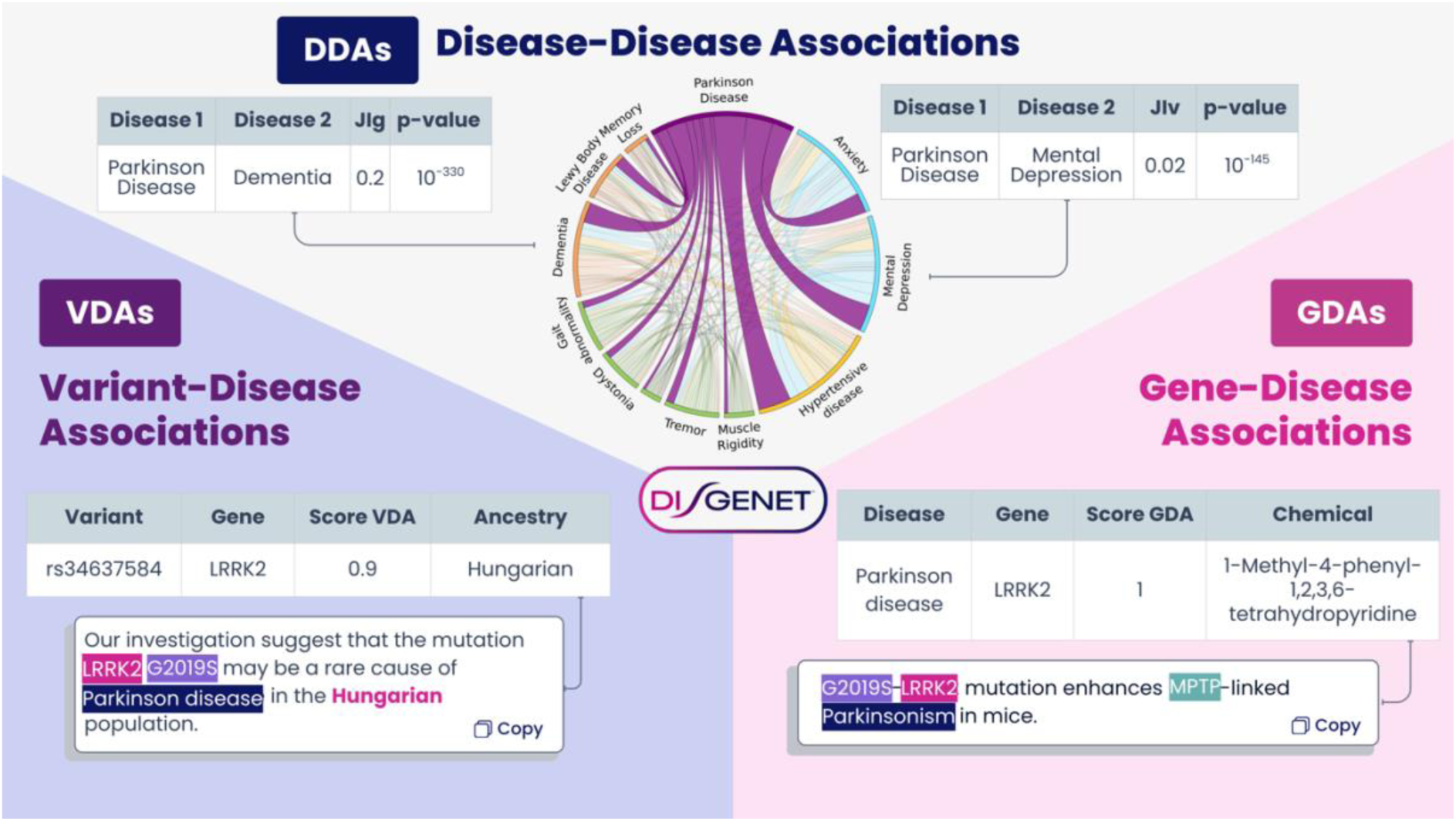
The DISGENET data model. The three main association types represented in DISGENET are the Gene–Disease Associations (GDAs), that combine multiple sources of evidence to link genes to human diseases, the Variant–Disease Associations (VDAs), which capture clinical and functional information connecting genetic variants to disease phenotypes, and the Disease–Disease Associations (DDAs) reflect shared molecular mechanisms, comorbidities, or phenotypic overlaps between diseases. For brevity, only some annotations and metrics are represented in the figure.

**Figure 2:**
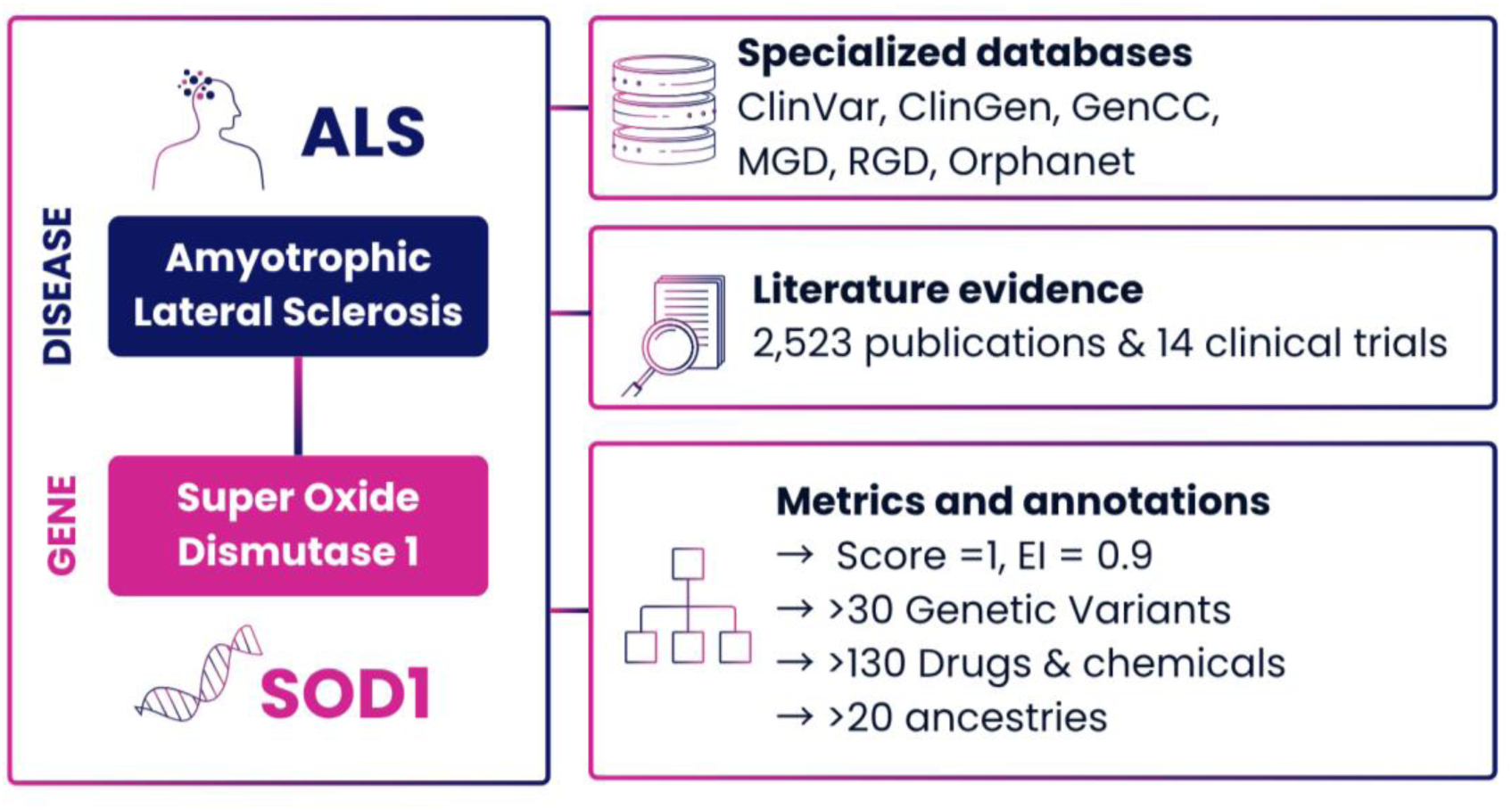
Conceptual framework for establishing a gene-disease association. The example of the superoxide dismutase 1 (SOD1) gene and Amyotrophic Lateral Sclerosis (ALS) illustrates the integration of specialized databases, literature evidence, and various metrics/annotations. For brevity, only some annotations and metrics are represented in the figure.

GDAs in DISGENET are extracted from a wide range of sources, systematically categorized to reflect their origin and nature. Curated GDAs are sourced from specialized databases such as UniProt, ClinVar, Orphanet, PsyGeNET, ClinGen, GenCC, MGD, and RGD, ensuring high-quality, expert-reviewed associations. Inferred GDAs are computationally derived by integrating data from the Human Phenotype Ontology (HPO), where gene-phenotype and phenotype-disease relationships are combined to infer GDAs, and from resources like the GWAS Catalog, PheWAS Catalog, and biobank datasets. Additionally, GDAs are extracted from the scientific literature and clinical trial records using automated text mining to capture the most recent findings and associations currently under investigation.

Figure 2 illustrates the comprehensive Gene-Disease Association (GDA) concept using the well-established link between the SOD1 gene and Amyotrophic Lateral Sclerosis (ALS). This association has a DISGENET score of 1.0 and an Evidence Index of 0.93, reflecting a robust body of evidence accumulated between 1993 and 2025. This includes over 2,500 scientific publications, 14 clinical trials examining protein levels and variants, and data from specialized databases such as ClinGen, GenCC, Orphanet, ClinVar, RGD, and MGD. Contextual information can provide key data to assess population relevance (across more than 20 ancestries and 39 genetic variants), generate mechanistic insights (DISGENET association type, presence of LoF or GoF mutations), and assess pharmacogenomic and toxicogenomic context (across over 130 drugs and chemicals, including Riluzole, Tofersen, and Arimoclomol). By integrating this multifaceted evidence, the GDA concept enables users to assess the maturity and significance of a gene-disease link, moving beyond a simple pairing to a multi-dimensional understanding. An illustration of how the DISGENET score is calculated is provided in Figure 3.

**Figure 3:**
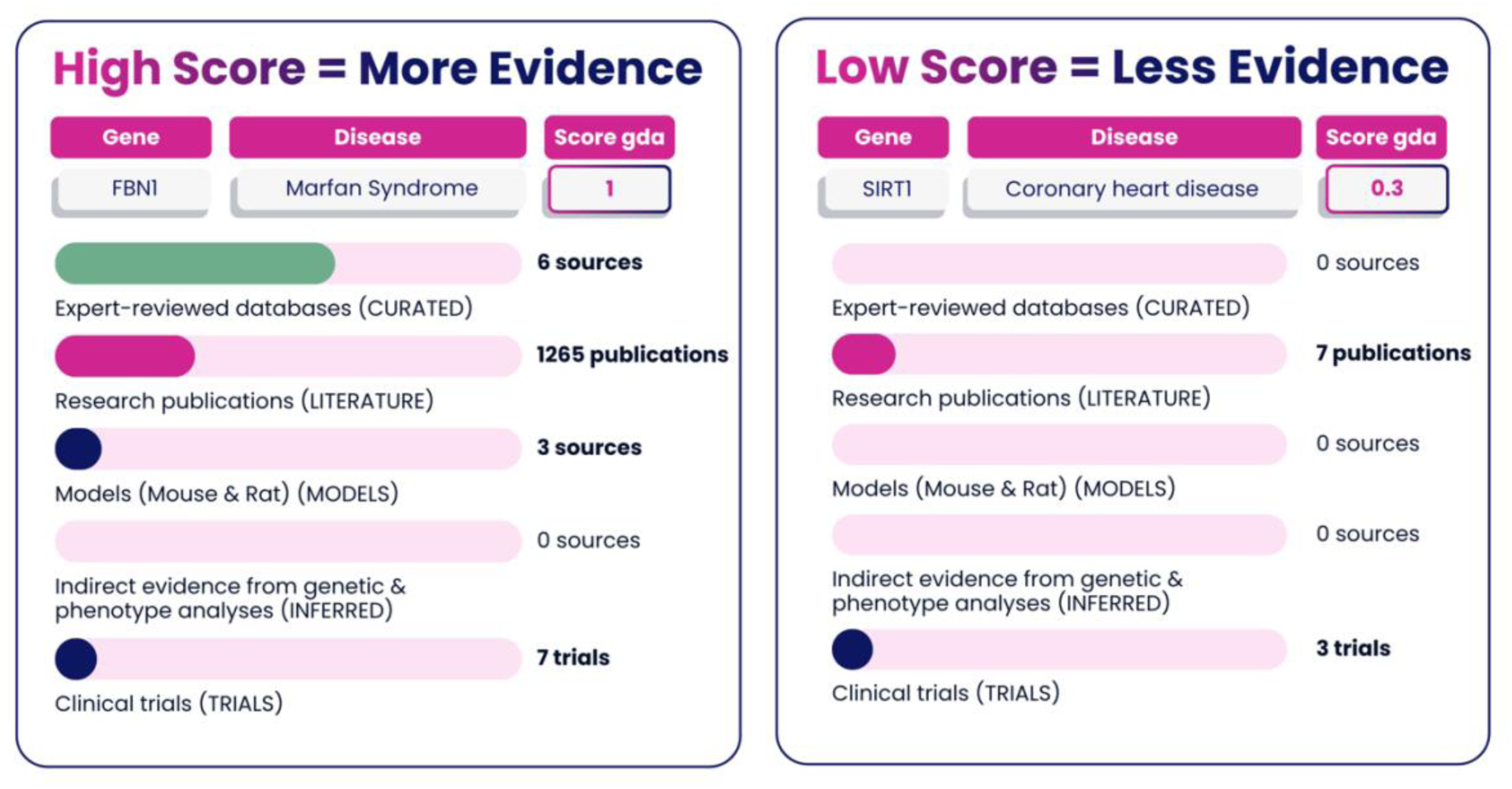
Illustration of the DISGENET score calculation. The figure compares a high-scoring, well-supported gene-disease association (FBN1-Marfan Syndrome, Score 1.0), supported by abundant evidence across multiple categories, with a lower-scoring association (SIRT1-Coronary heart disease, Score 0.3), supported by less evidence.

The variant-disease association (VDA) is the analogous concept for integrating evidence for genomic variants associated with diseases. Curated VDAs are obtained from specialized databases such as UniProt/SwissProt, ClinVar, the NHGRI-EBI GWAS Catalog, and the PheWAS Catalog, ensuring high-confidence annotations. Text Mining Human captures VDAs extracted from the scientific literature using text-mining approaches applied to human studies. Biobank data is a significant recent enhancement in DISGENET, expanding variant-disease associations with the integration of population-scale GWAS studies from FinnGen and the UK Biobank. These biobank-driven resources significantly enhance DISGENET coverage of population-scale genetic associations, providing a more granular view of the genetic architecture of diseases across large, well-characterized cohorts.

A unique feature of DISGENET is its comprehensive catalog of Disease-Disease Associations (DDAs), computationally inferred from shared genes and genetic variants. These associations are annotated with semantic similarity metrics computed over the Unified Medical Language System (UMLS) graph structure, and with semantic relations from the UMLS Metathesaurus, such as ’has_manifestation’ or ’is_complication_of’, to provide deeper mechanistic and clinical context. The DDA dataset can be used to: 1) explore the relationships among diseases, including disease comorbidities; 2) explore the links between diseases and their corresponding symptoms, signs, and phenotypes; or 3) identify genes or variants underlying disease associations. This information can be leveraged to support the drug R&D pipeline at multiple points. For instance, to develop new therapies for comorbid conditions, to stratify patients based on distinct disease profiles, or to identify biomarkers that overlap across multiple disease processes.

### Fine-grained Annotations with State-of-the-art Natural Language Processing

A distinctive feature of DISGENET is its ability to incorporate key contextual information into genotype-phenotype associations and to gather the most recent findings from the scientific literature. In fact, text-mined data accounts for around 70% of the database’s content, enabling DISGENET to capture emerging scientific discoveries, including those not yet curated by specialized resources. Moreover, this data enriches the associations collated by specialized resources with additional publication evidence, providing detailed provenance and broader literature support.

This extensive literature coverage is enabled by our proprietary Natural Language Processing (NLP) pipeline, which has continuously evolved to incorporate state-of-the-art biomedical text-mining methodologies. Designed for high accuracy^3^ and scalability, it systematically extracts and standardizes diverse biomedical entities and their relationships from the scientific literature. However, the rise of large language models (LLMs) has introduced both opportunities and challenges for biomedical text mining. Despite the widespread use of LLMs for identifying and extracting relevant data from biomedical texts, the complexities of finding, integrating, and harmonizing this disparate information remain a cumbersome and expensive process, posing a significant barrier to research. Generative AI-based approaches excel in reasoning-related tasks, such as question answering. However, they still perform worse than traditional NLP methods, including fine-tuned language models, for key information extraction from biomedical texts (8). In addition, hallucinations and their non-reproducible behavior limit their use for the standardization of NLP and data extraction procedures, not to mention their high computational cost and energy demands.

Our approach to NLP combines a variety of techniques (pattern matching, embeddings, supervised classification using traditional ML, fine-tuned Language Models, and LLMs) to efficiently identify GDAs, VDAs, and the effects of chemicals and drugs. It also captures crucial contextual information, such as how the gene (or variant) is linked to the phenotype (standardized with the DISGENET Association Type ontology), which population was assessed, the experimental model used, and whether loss-of-function (LoF) or gain-of-function (GoF) mutations influence the phenotype, among other data facets. These fine-grained annotations enable users to contextualize genotype-phenotype associations, facilitating interpretation and data-driven decision making. Crucially, references from the original publication and databases are provided for every piece of information, ensuring full traceability and allowing users to inspect the primary evidence directly.

### The DISGENET Suite of Tools

The DISGENET platform provides a comprehensive suite of tools designed to meet the needs of users with diverse backgrounds and skill sets, ensuring broad accessibility to its expanded database. This user-centric approach is reflected in the variety of access points available, including the open-source software solutions, the R package disgenet2r, and the DISGENET Cytoscape App, which facilitate data integration and analysis. Furthermore, the introduction of an AI Assistant for natural language queries highlights our commitment to making genotype–phenotype data more accessible and actionable for the entire scientific community, from wet-lab biologists to computational researchers.

The DISGENET web platform (https://www.disgenet.com/) features a redesigned interface that offers intuitive navigation, powerful search and filtering, and enhanced visualizations. It supports flexible queries across all entity types using free text or standard identifiers, along with advanced data filtering, download, and visualization capabilities through heatmaps, networks, and biological pathway views. Analytical functionalities, such as Disease Enrichment Analysis, are integrated into the platform and also accessible via the REST API, disgenet2r package, and DISGENET Cytoscape App. In addition, the platform offers comprehensive online documentation for input fields, clarifying the meaning of metrics and annotations, and guiding users throughout the data exploration process.

For computational biologists and data scientists, DISGENET offers Programmatic Access via a REST API (https://api.disgenet.com/doc/swagger). This allows for seamless integration of DISGENET data into custom analytical workflows and pipelines, facilitating large-scale data processing and automated analyses. In addition, it streamlines the integration of DISGENET data with other datasets, including omics and clinical data, for multimodal data analysis.

The R package disgenet2r (https://gitlab.com/medbio/disgenet2r) provides a convenient, familiar environment for researchers accustomed to the R statistical programming language. This package enables efficient analysis and manipulation of DISGENET data directly within the R ecosystem.

The DISGENET Cytoscape App (https://apps.cytoscape.org/apps/disgenetapp) is designed for researchers interested in network-based analysis. This application enables visualization and exploration of complex genotype-phenotype networks, providing an intuitive graphical representation of relationships in the DISGENET database. The App features an API automation module that enables programmatically executing various functionalities using R and Python, supporting the development of reproducible, scalable analysis workflows based on DISGENET data.

A new tool that facilitates access to DISGENET is the AI Assistant (https://disgenet.com/Assistant), which leverages agentic AI to interrogate the REST API. This tool empowers users to query the database in natural language, automatically translating plain-language questions into structured API queries. By doing so, and by summarizing key results in an accessible format, the DISGENET AI assistant significantly lowers the barrier for users without programming expertise or specific knowledge of the platform’s data model.

Finally, general documentation about the platform is available at the web interface and in the Help Center (https://support.disgenet.com/support/home).

### Key Advances

The new DISGENET platform introduces substantial scientific and technical improvements over its previous academic version (DisGeNET v7.0) and represents a significant leap forward in capabilities and utility.

The key advances in the new DISGENET platform are multifaceted:

1. **Increased Frequency of Data Updates:** The database is updated quarterly, ensuring that its content remains current with the latest findings in basic and clinical research. This enhanced update frequency significantly improves the timeliness and relevance of the data available to researchers.
2. **Expanded Data Coverage:** The platform offers a substantial increase in data coverage across genes, variants, and their associations with diseases. Specifically, the current release (v25.3) includes a **1,183%** increase in variant–disease associations (VDAs), a **753%** increase in variants, a **78%** increase in gene–disease associations (GDAs), a **40%** increase in genes, and a **27%** increase in diseases and phenotypes.
3. **New Data Types:** The integration of novel data types, including over 12,000 chemical and drug associations, contextualizes genotype-phenotype associations within a pharmacological and toxicological framework and adds the environmental factor dimension to complex diseases.
4. **New Data Sources**: The new platform features data from PHEWAS studies and from GWAS performed on biobanks (FinnGen and UK Biobank). It also includes GDAs extracted from Clinical Trials and from the Gene Curation Coalition (GenCC) Database.
5. **New Annotations**: GDAs and VDAs in DISGENET are annotated with ancestry information (either provided by the original source or extracted from the scientific publication), chemicals and drugs and their effects (therapeutic, toxic), VDAs are annotated as gain-of-function and loss-of-function, and genes are mapped to biological pathways and the protein target ontology.
6. **Improved accessibility:** The suite of tools, including the web platform, REST API, R package, and Cytoscape App, has been comprehensively updated to support new features and handle increased data volume, ensuring a seamless user experience and enhanced analytical capabilities. Moreover, we continue to support open-source bioinformatic software by maintaining the DISGENET Cytoscape App and the disgenet2r R package. New features are continuously incorporated into the suite of tools, such as advanced search capabilities, configurable filters, and new analytical functions, such as the disease enrichment analysis tool. A new tool has been added—an AI assistant that streamlines interactions with the database.
7. **New Sustainability Strategy:** Implementing a freemium model ensures the platform’s long-term viability and continuous updates, a critical factor for maintaining a dynamic bioinformatics resource.

### Sustainability model and community impact

A significant development in the new DISGENET platform is the implementation of a novel sustainability strategy based on a freemium model. This operational model is designed to ensure the long-term viability and continuous evolution of the resource. Under this model, academic users can obtain free subscription licenses following an application process, granting them full access to the database and open-source software tools. Concurrently, for-profit organizations can subscribe to the platform and leverage DISGENET for their research and development needs. This dual-tiered approach allows the platform to generate revenue from commercial partnerships, which in turn fund ongoing development and infrastructure maintenance. This strategic decision, implemented in 2020, addresses a common challenge faced by publicly funded bioinformatics resources, ensuring continuous updates and support, thereby guaranteeing the resource’s enduring utility and reliability in a dynamic research landscape.

Its extensive data integration and analytical capabilities have established DISGENET as a key resource for researchers worldwide for more than 15 years. The platform has been increasingly cited in scientific literature for its use in research and education^4^, including the investigation of disease mechanisms, drug discovery and development, clinical genomics, and the development of new AI approaches, among many other research areas (9–16). In addition, DISGENET data has been integrated into numerous third-party resources and software packages and used as a foundational dataset for various machine learning and AI tools (13,17–20).

As of October 2025, the platform’s adoption is demonstrated by the number of registered users (more than 16 K), monthly web visits (more than 10 K), bibliographic citations (more than 8 K scientific publications), and commercial subscriptions that sustain the resource.

### Applications across Research, Pharma, and Clinical Genomics

The availability of reliable, comprehensive human genotype–phenotype association data is fundamental across a wide range of research domains, underpinning key activities in therapeutic R&D and clinical genomics. DISGENET serves as a versatile platform for both basic and translational research, transforming genomic knowledge into actionable insights for academic researchers, the pharmaceutical industry, and health care organizations. The breadth and depth of its data, combined with detailed provenance and standardized annotations, make DISGENET a powerful and trustworthy resource for a wide range of biomedical applications. Furthermore, its suite of tools—including the REST API, disgenet2r package, and DISGENET Cytoscape App —facilitates the integration of third-party or user-provided data with DISGENET content, enabling comprehensive multi-modal analyses that are essential for translational research and drug development.

DISGENET is a versatile platform that supports several key points of the therapeutic R&D pipeline, from pre-clinical to clinical stages (Figure 4). DISGENET is not only effective for target identification and prioritization (21–24) but also for target de-risking (25), biomarker discovery (26,27), patient stratification (28,29), assessment of white space opportunities, drug repurposing (30), and indication expansion. Thanks to its extensive data coverage — over 2 million Gene-Disease Associations (GDAs), 4 million Variant-Disease Associations (VDAs), and over 20 million DDAs — it can streamline the identification of both drug targets and biomarkers based on pre-clinical and clinical evidence. It enables discovery across all therapeutic areas thanks to its broad phenotypic spectrum: Mendelian and Complex disorders, phenotypic traits and symptoms, and phenotypes observed as drug adverse effects. The extensive disease profiles for genes and variants enable the identification of potential safety issues of candidate drugs in early phases, as well as the assessment of biomarker specificity in disease contexts. The selection and prioritization of targets and biomarkers alike can be expedited by the variety of metrics and annotations that DISGENET offers, such as:

- The **DISGENET Score** enables the selection of associations based on the strength of evidence from literature and databases, ranging from well-supported to emerging findings (see Figure 3 for an example of the DISGENET score calculation).
- **The DSI (Disease Specificity Index) and the DPI (Disease Pleiotropy Index)** are used to identify pleiotropic genes and variants.
- **The Evidence Index** to identify associations with contradictory evidence
- **Variant allele frequency and GWAS summary statistics** to support target nomination
- **The DISGENET Association Type**, **LoF and GoF** annotations, **variant consequence types**, and **deleteriousness scores** help the user understand how the gene (or variant) is linked with the phenotype and gain mechanistic insights.
- Information about **ancestry** provides insights for population-specific assessment of targets and biomarkers, along with **variant allele frequency** and **disease prevalence**, addressing pharmaceutical needs for developing therapies tailored to genetic diversity.
- GDAs and VDAs are annotated with **drugs and their effects**, enabling the assessment of pharmacogenomic associations.

**Figure 4:**
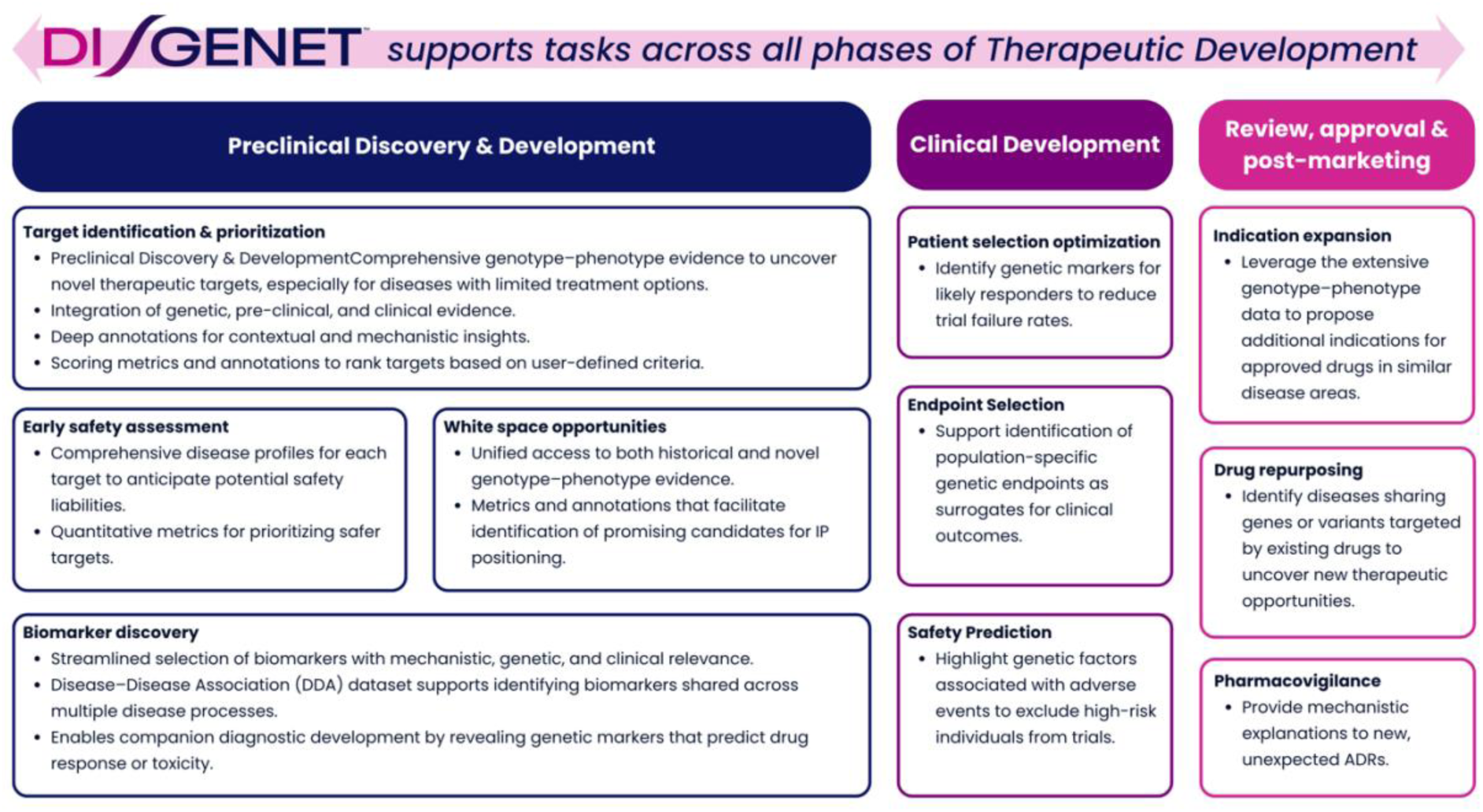
DISGENET applications across the therapeutic R&D pipeline. The flowchart illustrates platform utility from target identification through clinical applications, highlighting key features supporting each phase: (A) Pre-clinical discovery and development, (B) Clinical Development, (C) Post-marketing applications.

Last but not least, data standardization facilitates the combined analysis of DISGENET data with other datasets, supporting integrative analysis across multiple steps of the therapeutic R&D pipeline.

### DISGENET in Clinical Genomics

In recent years, Whole Exome Sequencing and Whole Genome Sequencing (WGS) have been increasingly adopted in clinical settings. In particular, WGS offers advantages over other approaches (by enabling the detection of a broader range of variants and providing untargeted coverage), but introduces significant challenges (the interpretation of the clinical relevance of variants in non-coding and intergenic regions) (31,32). Notwithstanding these challenges, the contribution of non-coding variants to Mendelian disease is increasingly recognized (33). Therefore, laboratories employing WGS should ensure that their pipelines can adequately analyze non-coding variation, including its annotation with disease-relevant information. Notably, non-coding variants are underrepresented in clinical variant databases such as ClinVar (34). As a result, complementary databases that capture the full spectrum of genomic variation should be integrated into variant interpretation workflows to realize the potential of WGS fully. In this context, the distinctive features of DISGENET make it a powerful resource for clinical genomics. These include the integration of multiple sources of evidence, support for multi-gene analyses, extensive coverage of variants in non-coding regions, ancestry-specific insights, and a suite of metrics for advanced variant interpretation. Furthermore, DISGENET combines clinically validated data (from resources like ClinGen and GenCC) with emerging evidence on genotype–phenotype associations, providing a comprehensive framework for variant assessment. Box 1 illustrates how DISGENET supports gene and variant interpretation in clinical genomics.

Several companies, research centers, and health care organizations use DISGENET for NGS data analysis and precision medicine applications. For instance, DISGENET has been used to enhance bioinformatic pipelines for rare disease genomic analysis and clinical diagnosis (35–39). By leveraging DISGENET’s capability to uncover novel genotype-phenotype associations, clinical diagnosis support tools integrating DISGENET increased the diagnostic yield for rare disease patients and provided key information for clinical decision-making, thereby improving patient management (39). In a study focused on DNA methylation in patients with Cri du Chat syndrome, DISGENET identified connections between genes with promoter DNA methylation changes and disease phenotypes, suggesting a causal role for DNA methylation in this understudied rare disease (36). In a recent WES study of individuals from the Arabian Peninsula, DISGENET helped identify the clinical relevance of novel variants in this cohort, including 185 UTR variants and 76 nonsynonymous variants associated with neurologic and congenital diseases (37). Finally, DISGENET metrics have been used to develop a scoring approach for prioritizing rare autism spectrum disorder candidate variants from whole-exome sequencing (35).

##### Box 1

###### Extensive data coverage

- 4 Million variant-disease associations (VDAs) identified from pre-clinical and clinical studies and GWAS from population-based biobanks, in coding and non-coding regions of the genome, to support the interpretation of WGS data
- 2 Million gene-disease associations (GDAs) to support the interpretation of sequencing data and the design of gene panels
- Variants from all chromosomes, including the mitochondrial genome
- Broad coverage of the phenotype spectrum, including Mendelian and complex disorders, disease traits, and phenotypes

###### Advanced Metrics

- DISGENET Score enables selection of associations based on the strength of evidence from literature and databases, ranging from well-supported to emerging findings.
- DSI & DPI metrics to filter variants specific to a particular phenotype, helping in differential diagnosis.
- Evidence Index to identify associations with contradictory evidence (scoring metric indicating the consistency of supporting evidence)
- Allele frequency from large-scale studies to filter variants for further processing

###### Fine-grained annotations to support comprehensive insights

- ACMG classification for variants and GenCC gene validity levels to enable interpretation according to clinical standards
- DISGENET Association Type to support assessment of gene and variant impact on the phenotype
- Standardized ancestry information for population-specific interpretation
- Functional impact of variants to guide clinical decision-making, including LoF/GoF annotations, variant consequence types, and deleteriousness scores
- Pharmacogenomic Information: VDAs and GDAs are annotated with drugs, enabling the discovery of pharmacogenomic associations
- Evidence Provenance enables the tracking of the source of evidence, whether it comes from specialized databases (e.g., GenCC, Orphanet), literature-mined, GWAS, or Clinical Trials
- Disease and traits are standardized with popular ontologies, enabling seamless integration with clinical and research data

## Conclusion

DISGENET has evolved from a pioneering academic resource into an essential infrastructure supporting basic and translational research. By addressing critical challenges in data fragmentation and accessibility, the platform enables users to transform complex genomic information into actionable knowledge for drug discovery, clinical diagnostics, and precision medicine. Future developments will focus on expanding population-specific annotations, integrating multi-omics data, and enhancing AI-driven tools for automated knowledge discovery and data-driven decision support. The platform’s sustainable model ensures continued growth and adaptation to emerging scientific needs, positioning DISGENET as a cornerstone resource for the next generation of biomedical discoveries.

## Author contributions

Conceptualization: LIF, JP

Data curation: LIF, JP, NR

Formal analysis: JP

Funding acquisition: LIF, FS

Project administration: LIF, FS

Software: JP, JC, NR, AG, AM, DS, MDB, MDSS, IR, JTZ

Validation: JP, LIF

Visualization: JP, SH

Writing – original draft: JP, LIF

Writing – review and editing: JP, LIF, SH, FS

## Footnotes

^1^https://elixir-europe.org/intranet/forms/rir-selection/sid432

^2^Compared to DisGeNET v7.0 (the latest academic version previously available at https://www.disgenet.org/)

^3^Our latest benchmarking exercise resulted in an average 0.98 F1 score for entity recognition and 0.92 F1 score for relation extraction

^4^How Harvard Medical School Uses DISGENET | Case Study

## References

1. Bauer-Mehren A, Rautschka M, Sanz F, Furlong LI. DisGeNET: a Cytoscape plugin to visualize, integrate, search and analyze gene–disease networks. Bioinformatics. 2010 Nov 15;26(22):2924–6.

2. Bauer-Mehren A, Bundschus M, Rautschka M, Mayer MA, Sanz F, Furlong LI. Gene-Disease Network Analysis Reveals Functional Modules in Mendelian, Complex and Environmental Diseases. Khanin R, editor. PLoS ONE. 2011 June 14;6(6):e20284.

3. Pinero J, Queralt-Rosinach N, Bravo A, Deu-Pons J, Bauer-Mehren A, Baron M, et al. DisGeNET: a discovery platform for the dynamical exploration of human diseases and their genes. Database. 2015 Apr 15;2015(0):bav028–bav028.

4. Queralt-Rosinach N, Piñero J, Bravo À, Sanz F, Furlong LI. DisGeNET-RDF: harnessing the innovative power of the Semantic Web to explore the genetic basis of diseases. Bioinformatics. 2016 July 15;32(14):2236–8.

5. Piñero J, Bravo À, Queralt-Rosinach N, Gutiérrez-Sacristán A, Deu-Pons J, Centeno E, et al. DisGeNET: a comprehensive platform integrating information on human disease-associated genes and variants. Nucleic Acids Res. 2017 Jan 4;45(D1):D833–9.

6. Piñero J, Ramírez-Anguita JM, Saüch-Pitarch J, Ronzano F, Centeno E, Sanz F, et al. The DisGeNET knowledge platform for disease genomics: 2019 update. Nucleic Acids Res. 2019 Nov 4;gkz1021.

7. Piñero J, Saüch J, Sanz F, Furlong LI. The DisGeNET cytoscape app: Exploring and visualizing disease genomics data. Comput Struct Biotechnol J. 2021;19:2960–7.

8. Chen Q, Hu Y, Peng X, Xie Q, Jin Q, Gilson A, et al. Benchmarking large language models for biomedical natural language processing applications and recommendations. Nat Commun. 2025 Apr 6;16(1):3280.

9. Shen Y, Timsina J, Heo G, Beric A, Ali M, Wang C, et al. CSF proteomics identifies early changes in autosomal dominant Alzheimer’s disease. Cell. 2024 Oct;187(22):6309–6326.e15.

10. Jiang F, Hu SY, Tian W, Wang NN, Yang N, Dong SS, et al. A landscape of gene expression regulation for synovium in arthritis. Nat Commun. 2024 Feb 15;15(1):1409.

11. Kozawa S, Tejima K, Takagi S, Kuroda M, Nogami-Itoh M, Kitamura H, et al. Latent inter-organ mechanism of idiopathic pulmonary fibrosis unveiled by a generative computational approach. Sci Rep. 2023 Dec 11;13(1):21981.

12. Pinheiro-de-Sousa I, Fonseca-Alaniz MH, Giudice G, Valadão IC, Modestia SM, Mattioli SV, et al. Integrated systems biology approach identifies gene targets for endothelial dysfunction. Mol Syst Biol. 2023 Dec 6;19(12):e11462.

13. Queen O, Huang Y, Calef R, Giunchiglia V, Chen T, Dasoulas G, et al. ProCyon: A multimodal foundation model for protein phenotypes [Internet]. Bioinformatics; 2024 [cited 2025 June 7]. Available from: http://biorxiv.org/lookup/doi/10.1101/2024.12.10.627665

14. Clément AE, Merdrignac C, Puiggros SR, Sévère D, Brionne A, Lafond T, et al. Parent-of-origin regulation by maternal auts2 shapes neurodevelopment and behavior in fish. Genome Biol. 2025 May 9;26(1):125.

15. Zong N, Chowdhury S, Zhou S, Rajaganapathy S, Yu Y, Wang L, et al. Advancing efficacy prediction for electronic health records based emulated trials in repurposing heart failure therapies. Npj Digit Med. 2025 May 24;8(1):306.

16. Gonzalez-Lozano MA, Schmid EW, Miguel Whelan E, Jiang Y, Paulo JA, Walter JC, et al. EndoMAP.v1 charts the structural landscape of human early endosome complexes. Nature [Internet]. 2025 May 28 [cited 2025 June 7]; Available from: https://www.nature.com/articles/s41586-025-09059-y

17. Huang K, Fu T, Gao W, Zhao Y, Roohani Y, Leskovec J, et al. Artificial intelligence foundation for therapeutic science. Nat Chem Biol. 2022 Oct;18(10):1033–6.

18. Huang K, Zhang S, Wang H, Qu Y, Lu Y, Roohani Y, et al. Biomni: A General-Purpose Biomedical AI Agent [Internet]. Bioinformatics; 2025 [cited 2025 Oct 29]. Available from: 10.1101/2025.05.30.656746

19. Bonner S, Barrett IP, Ye C, Swiers R, Engkvist O, Bender A, et al. A review of biomedical datasets relating to drug discovery: a knowledge graph perspective. Brief Bioinform. 2022 Nov 19;23(6):bbac404.

20. Chaves JMZ, Wang E, Tu T, Vaishnav ED, Lee B, Mahdavi SS, et al. Tx-LLM: A Large Language Model for Therapeutics [Internet]. arXiv; 2024 [cited 2025 Oct 29]. Available from: http://arxiv.org/abs/2406.06316

21. Blaudin De Thé FX, Baudier C, Andrade Pereira R, Lefebvre C, Moingeon P. Transforming drug discovery with a high-throughput AI-powered platform: A 5-year experience with Patrimony. Drug Discov Today. 2023 Nov;28(11):103772.

22. Zanardi A, Nardini I, Raia S, Conti A, Ferrini B, D’Adamo P, et al. New orphan disease therapies from the proteome of industrial plasma processing waste- a treatment for aceruloplasminemia. Commun Biol. 2024 Jan 30;7(1):140.

23. Namba S, Iwata M, Nureki SI, Yuyama Otani N, Yamanishi Y. Therapeutic target prediction for orphan diseases integrating genome-wide and transcriptome-wide association studies. Nat Commun. 2025 Apr 18;16(1):3355.

24. Breimann S, Kamp F, Basset G, Abou-Ajram C, Güner G, Yanagida K, et al. Charting γ-secretase substrates by explainable AI. Nat Commun. 2025 July 1;16(1):5428.

25. Carss KJ, Deaton AM, Del Rio-Espinola A, Diogo D, Fielden M, Kulkarni DA, et al. Using human genetics to improve safety assessment of therapeutics. Nat Rev Drug Discov. 2023 Feb;22(2):145–62.

26. Pistilli B, Mosele F, Corcos N, Pierotti L, Pradat Y, Le Bescond L, et al. Patritumab deruxtecan in HR+HER2− advanced breast cancer: a phase 2 trial. Nat Med. 2025 Oct;31(10):3492–503.

27. Mao G, Cong W. Identification and experimental validation of biomarkers related to mitochondrial and programmed cell death in obsessive-compulsive disorder. Sci Rep. 2025 Aug 30;15(1):31971.

28. Hernández-Lorenzo L, García-Gutiérrez F, Solbas-Casajús A, Corrochano S, Matías-Guiu JA, Ayala JL. Genetic-based patient stratification in Alzheimer’s disease. Sci Rep. 2024 Apr 30;14(1):9970.

29. Velz J, Freudenmann LK, Medici G, Dubbelaar M, Mohme M, Ghasemi DR, et al. Mapping naturally presented T cell antigens in medulloblastoma based on integrative multi-omics. Nat Commun. 2025 Feb 4;16(1):1364.

30. Tanoli Z, Fernández-Torras A, Özcan UO, Kushnir A, Nader KM, Gadiya Y, et al. Computational drug repurposing: approaches, evaluation of in silico resources and case studies. Nat Rev Drug Discov [Internet]. 2025 Mar 18 [cited 2025 June 7]; Available from: https://www.nature.com/articles/s41573-025-01164-x

31. Bagger FO, Borgwardt L, Jespersen AS, Hansen AR, Bertelsen B, Kodama M, et al. Whole genome sequencing in clinical practice. BMC Med Genomics. 2024 Jan 29;17(1):39.

32. Austin-Tse CA, Jobanputra V, Perry DL, Bick D, Taft RJ, Venner E, et al. Best practices for the interpretation and reporting of clinical whole genome sequencing. Npj Genomic Med. 2022 Apr 8;7(1):27.

33. French JD, Edwards SL. The Role of Noncoding Variants in Heritable Disease. Trends Genet. 2020 Nov;36(11):880–91.

34. Ellingford JM, Ahn JW, Bagnall RD, Baralle D, Barton S, Campbell C, et al. Recommendations for clinical interpretation of variants found in non-coding regions of the genome. Genome Med. 2022 Dec;14(1):73.

35. Shil A, Arava N, Levi N, Levine L, Golan H, Meiri G, et al. An integrative scoring approach for prioritization of rare autism spectrum disorder candidate variants from whole exome sequencing data. Sci Rep. 2025 Apr 15;15(1):13024.

36. Holland P, Wildhagen M, Istre M, Reiakvam OM, Dahl JA, Søraas A. Cri du chat syndrome patients have DNA methylation changes in genes linked to symptoms of the disease. Clin Epigenetics. 2022 Dec;14(1):128.

37. Ferreira JC, Alshamali F, Pereira L, Fernandes V. Characterization of Arabian Peninsula whole exomes: Contributing to the catalogue of human diversity. iScience. 2022 Nov;25(11):105336.

38. Núñez-Carpintero I, Rigau M, Bosio M, O’Connor E, Spendiff S, Azuma Y, et al. Rare disease research workflow using multilayer networks elucidates the molecular determinants of severity in Congenital Myasthenic Syndromes. Nat Commun. 2024 Feb 28;15(1):1227.

39. Chetta M, Tarsitano M, Rivieccio M, Oro M, Cammarota AL, De Marco M, et al. A Copernican revolution of multigenic analysis: A retrospective study on clinical exome sequencing in unclear genetic disorders. Comput Struct Biotechnol J. 2024 Dec;23:2615–22.

